# African Swine Fever Virus major capsid p72 Trimers function as a pH sensor during uncoating process of virus endocytosis

**DOI:** 10.1101/2023.06.09.544430

**Authors:** Kaiwen Meng, Yangnan Huyan, Qi Liu, Junyi Li, Ye Xiang, Geng Meng

**Affiliations:** College of Veterinary Medicine, China Agricultural University, 100193, Beijing, China; School of Medicine, Tsinghua University, 100084, Beijing, Chin

## Abstract

African swine fever virus (ASFV) belongs to nucleocytoplasmic large DNA viruses (NCLDVs), the only member of Asfarviridae. So far, it is revealed that the ASFV uncoating is a pH-dependent process undergone in late endosomes. But, the research on how pH affects capsid stability is limited, and which protein plays an essential role in pH sensing remains unknown. In this study, we identified the main component of the ASFV capsid -- major capsid protein p72, as a pH sensitive residue abundance protein, and it is speculated that the conformational change of the p72 trimer is possibly responsible for the ASFV uncoating process. To test this speculation, we obtained recombinant p72 trimers, treated with the acidic environment that simulated endosomes and displayed structural analysis. The results showed that the p72 trimer depolymerized at low pH. The depolymerization of trimers rationally explains the disassembly mechanism of the ASFV icosahedral capsid in endosomes.

## Introduction

The nucleocytoplasmic large DNA viruses (NCLDV) are a group of large double-strand DNA (dsDNA) viruses. Many of the NCLDVs, including African swine fever virus (ASFV), Iridovirus, Mimivirus, Paramecium bursaria chlorella Virus (PBCV), Faustovirus,(FV) have icosahedral or close-to-icosahedral capsid symmetry assembled from trimeric major capsid protein (MCP) with typical double jelly-roll (DJR) fold. As the only known member of icosahedral NCLDVs that infect mammals, ASFV belongs to the Asfarviridae, and can cause acute hemorrhagic fever in pigs and wild boars. It is a highly contagious and lethal virus (1-4). At present, ASFV has not been completely eliminated in China and neighboring countries.

Similar to other NCLDVs, the ASFV is multi-layered. The ASFV particle is composed of an exterior membrane envelope, an icosahedral capsid (T=277), an inner membrane, an icosahedral core-shell (T=17) and an inner core containing the genome (5). Therefore, in the uncoating process, the viral capsid should be disassembled to release the enveloped core shell and further release the genome. However, as a key step in the replication cycle, the uncoating mechanism of ASFV, as well as NCLDVs and other dsDNA viruses, is very limited so far.

Previous studies revealed that ASFV enters host cells through the endosomal pathway (6-9). There is a pH gradient between the vesicles of the endosome system. The pH in the early endosomes is close to 7, and the pH is around 5-6 in the late endosome. Electron microscopic observation of ASFV entering macrophages showed that the density of ASFV capsid is partially lost during the virus transportation from early endosome to late endosome (7). And it has been verified that the ASFV capsid disruption relies on endosomal acidification (7). Similar endosomal pathway/low pH-dependent uncoating processes have also been reported in other dsDNA viruses (10-12). However, the pH-sensing responding component of the ASFV capsid still remains unknown.

In this paper, by comparing the histidine component of all capsid proteins, we identified the major capsid protein p72 as a pH sensor. The p72 trimer is the main component of the outer capsid of virus particles. Therefore, it is reasonable to speculate that the pH-dependent conformational change of p72 trimer is likely to be the starting point of the uncoating process. Here, we successfully verified that the recombinant p72 could undergo depolymerization after being treated in acidic environment. The same depolymerization phenomenon may also happen on the p72 trimer in the particle transferred to the late endosome. This finding may help to understand the uncoating process of ASFV and other NCLDVs virus particles.

## Results

### The major capsid protein p72 is rich in histidine

Histidine is generally regarded as a class of pH-sensitive amino acids that mediate conformational changes in protein molecules (13-15). Thus, we determined and compared the histidine content within the structural proteins located on the capsid, in order to identify pH sensor. Based on previous proteomic and ASFV structural studies (5), the following proteins have been identified as localized in the capsid: p72, p49 (pB438L), H240R, pE120R and M1249L. The results showed that the content of histidine in p72, p49, H240R, pE120R and M1249L was 4.80% (31/646), 4.11% (18/438), 3.75% (9/240), 3.33% (4/120) and 2.08% (26/1249), respectively (Figure 1). Among them, p72 has the highest content of histidine, and is most likely to play a major role in capsid pH sensing.

**Figure 1.**
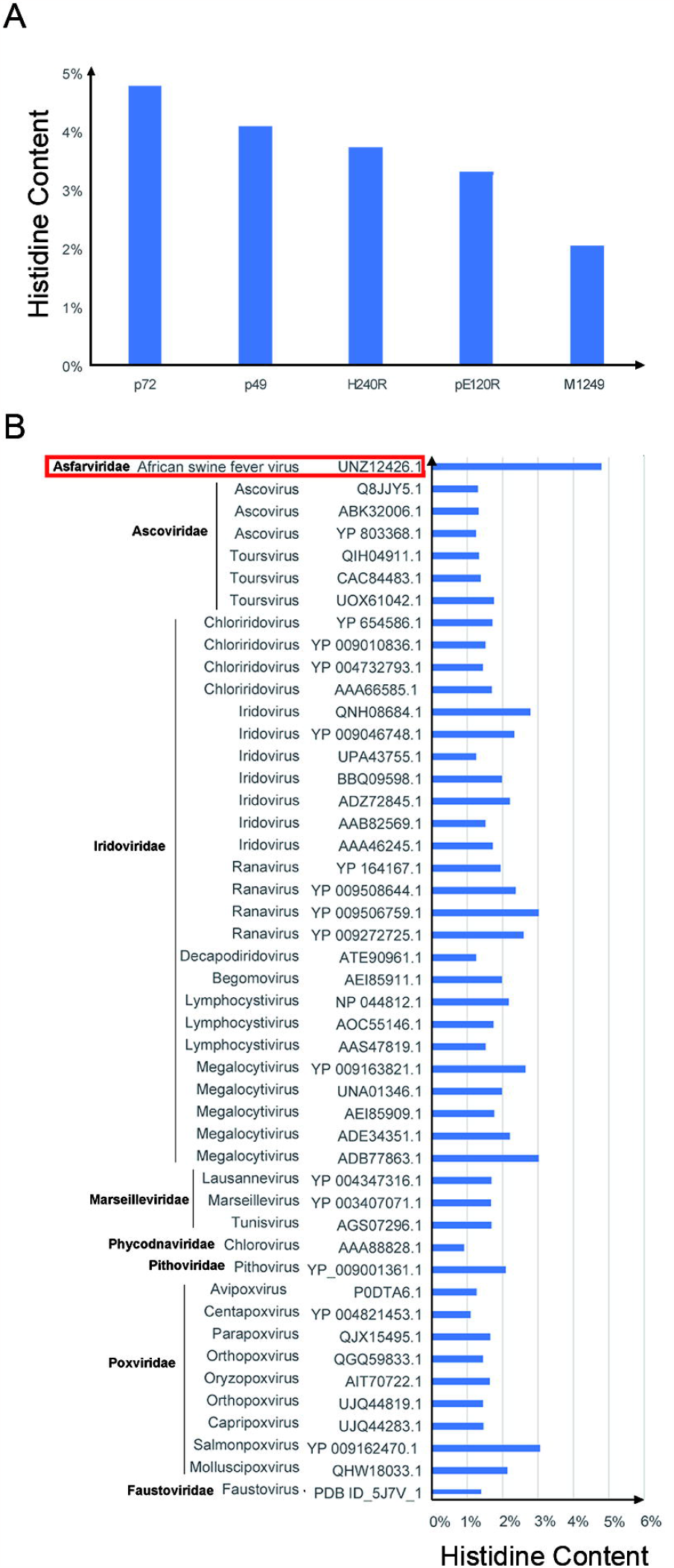
Histidine content in various NCLDVs capsid protein. (A) Analysis of the histidine content of structural proteins located in the ASFV capsid. (B) Comparison of histidine content in NCLDVs capsid protein.

The high histidine content is also a unique feature of ASFV p72 compared with other MCP of NCLDVs. We calculated the proportion of histidine in the MCP sequences of NCLDVs previously collected from NCBI (Figure 1B). The results showed that the histidine content of ASFV p72 is at least 1.5 folds of that in other NCLDVs MCP. Of note, other than ASFV, members of the family Iridoviridae had slightly higher average histidine content in MCP than other NCLDVs. The average histidine content of Iridoviridae MCP was 2.02%, while the average histidine content of the MCP of Ascoviridae, Marseilleviridae and Poxviridae was 1.39%, 1.68%, and 1.69%, respectively. Besides, the Iridoviridae is also the closest relative to Asfarviridae among the NCLDVs that could infect vertebrates (Figure 2B, Table S1).

**Figure 2.**
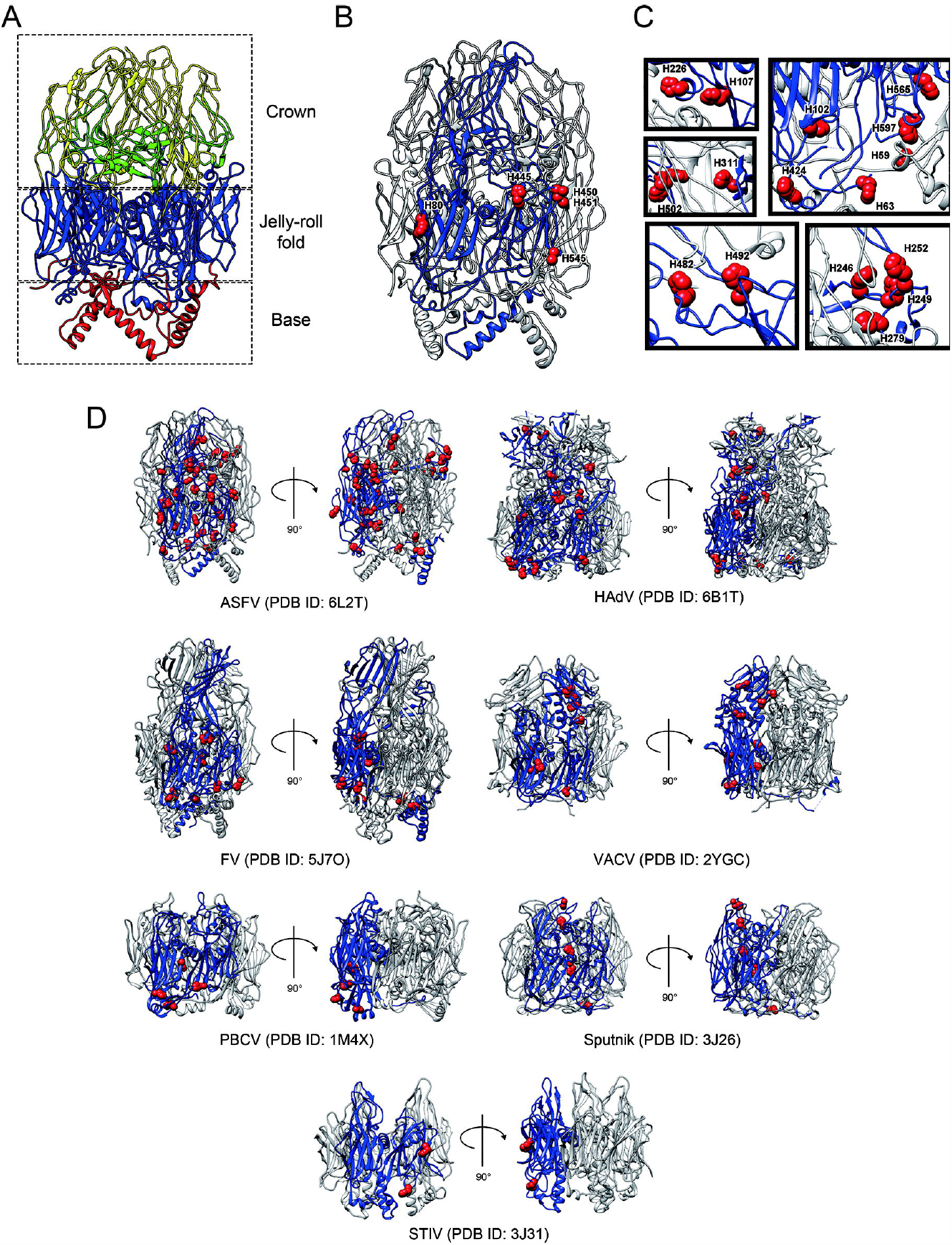
Histidine distribution in DJR protein. (A) Ribbon diagram shows the structure of p72 trimer. The base domain is colored in red, the jelly-roll fold is colored in blue, the crown domain is colored in yellow, and the F1-G1 loop is colored in green. (B-C) Histidines with side chains exposed on the surface of ASFV p72 monomers. One of the monomers in the trimer is shown in blue and the remaining two monomers are shown in grey. Histidines whose side chains are exposed on the surface of the p72 monomer are shown as red spheres, including histidines exposed at the inter-trimer contact interface (B) and histidines exposed at the inter-trimer contact interface (C). (D) Histidine distribution in capsid proteins of DJR. One of the monomers in the trimer is shown in blue and the remaining two monomers are shown in grey.

### Histidine in p72 is mostly localized on the interface within the trimer

Except for NCLDVs, double jelly-roll (DJR) fold is also a common feature of the MCP of several dsDNA viruses, including Adenoviridae, Lavidaviridae and Turriviridae. We collected and compared all published high-resolution structures of MCP with DJR fold, significant deviation in histidine distribution was observed (Table 1, Figure 2A). Of note, the histidines in ASFV p72 were distributed in both DJR domain and crown domain. And the histidine in the crown domain is mainly located in the F1-G1 loop. While in the Faustovirus (FV) MCP, whose structure is highly similar to p72, the histidine is only distributed in the DJR fold domain.

**Table 1.** Histidine content in DJR capsid protein

| Virus | Species | PDB ID | Samples | Method | Resolution ( Å ) | Content of histidine |
| --- | --- | --- | --- | --- | --- | --- |
| ASFV | *Asfarviridae* | 6KU9<sup>(26)</sup> | Recombinant p72 | Cryo-EM | 2.67 | 4.8% (31/646) |
| Human Adenovirus (HAdV) | *Adenoviridae* | 6B1T<sup>(27)</sup> | Viral particles cultured in vitro | Cryo-EM | 3.2 | 1.68% (16/952) |
| Faustovirus (FV) | *Faustoviridae* | 5J7O<sup>(22)</sup> | MCP extracted from whole virus | X-ray diffraction | 2.37 | 1.4% (9/644) |
| Vaccinia virus (VACV) | *Poxviridae* | 2YGC<sup>(28)</sup> | Recombinant D13 protein | X-ray diffraction | 3.02 | 1.45% (8/551) |
| Paramecium bursaria chlorella Virus (PBCV) | *Phycodnaviridae* | 1M4X<sup>(29)</sup> | p54 extracted from whole virus | X-ray diffraction | 2 | 0.97% (4/413) |
| Sputnik virophage | *Lavidaviridae* | 3J26<sup>(11)</sup> | Purified virions from infected tissues | Cryo-EM | 3.5 | 0.98% (5/508) |
| Sulfolobus turreted icosahedral virus (STIV) | *Turriviridae* | 3J31<sup>(30)</sup> | Recombinant MCPs and purified virions from infected tissues | Cryo-EM and X-ray diffraction | 4.5 | 0.58% (2/345) |

Further analysis based on high-resolution structural information showed that in p72, at least 21 histidines’ side chains are exposed on the surface of the monomer (Figure 2C-2D). Among them, 5 histidines in the DJR fold domain are located on the inter-trimer contact surface, including H80, H445, H450, H451 and H545 (Figure 2C). And the other 16 histidines are located on the intra-trimer contact surface between adjacent monomers (Figure 2D), including 9 histidines in the crown domain (H226, H246, H249, H252, H278, H311, H252, H278, H311, H482, H492 and H502), 3 histidines in the N-terminal jelly-roll (JR) domain (H102, H107, H424), 2 histidines in the C-terminal JR domain (H565, H597) and 2 histidines in the linker between Base domain and JR domain (H59, H63).

### The p72 particle size reduction after treated with low pH

According to different environments that the ASFV may expose to during viral invasion, three pHs were selected, including pH 5.5, pH 5 and pH 3. The buffer of pH 5.5 and pH 5 were used to mimic the acidic environment in late endosome and lysosome, respectively. As porcine can be infected by intake feeds polluted by ASFV, we prepared a buffer of pH 3 to test the effect of a stomach-similar environment on the stability of p72.

The recombinant trimer was prepared according to a previous report by Qi Liu *et al*.(16). According to their study, the full-length native p72 could only be folded correctly into trimer with the aid of B602L. Here, we verified the production of the co-expressed p72 and B602L by SDS-PAGE (Figure S2) and mass spectrometry (MS) (Figure S3).

Size exclusion chromatography analysis was performed. The results showed that the peak position of p72 after acid treatment moved backwards significantly compared with that of the untreated sample (Figure 3A). According to the standard curve of peak position and molecular weight previously established in the laboratory, the molecular weight corresponding to the peak position of untreated p72 (pH 8.5) was calculated to be about ∼200kDa. After being treated with pH 5.5 and pH 5, the theoretical molecular weight of samples becomes ∼150kDa. After being treated with pH 3, the theoretical molecular weights of samples were changed to ∼100kDa. These results indicated that the particle size of p72 sample decreased by 25% after being treated with Ph 5.5 and pH 5, and decreased by 50% at pH 3.

**Figure 3.**
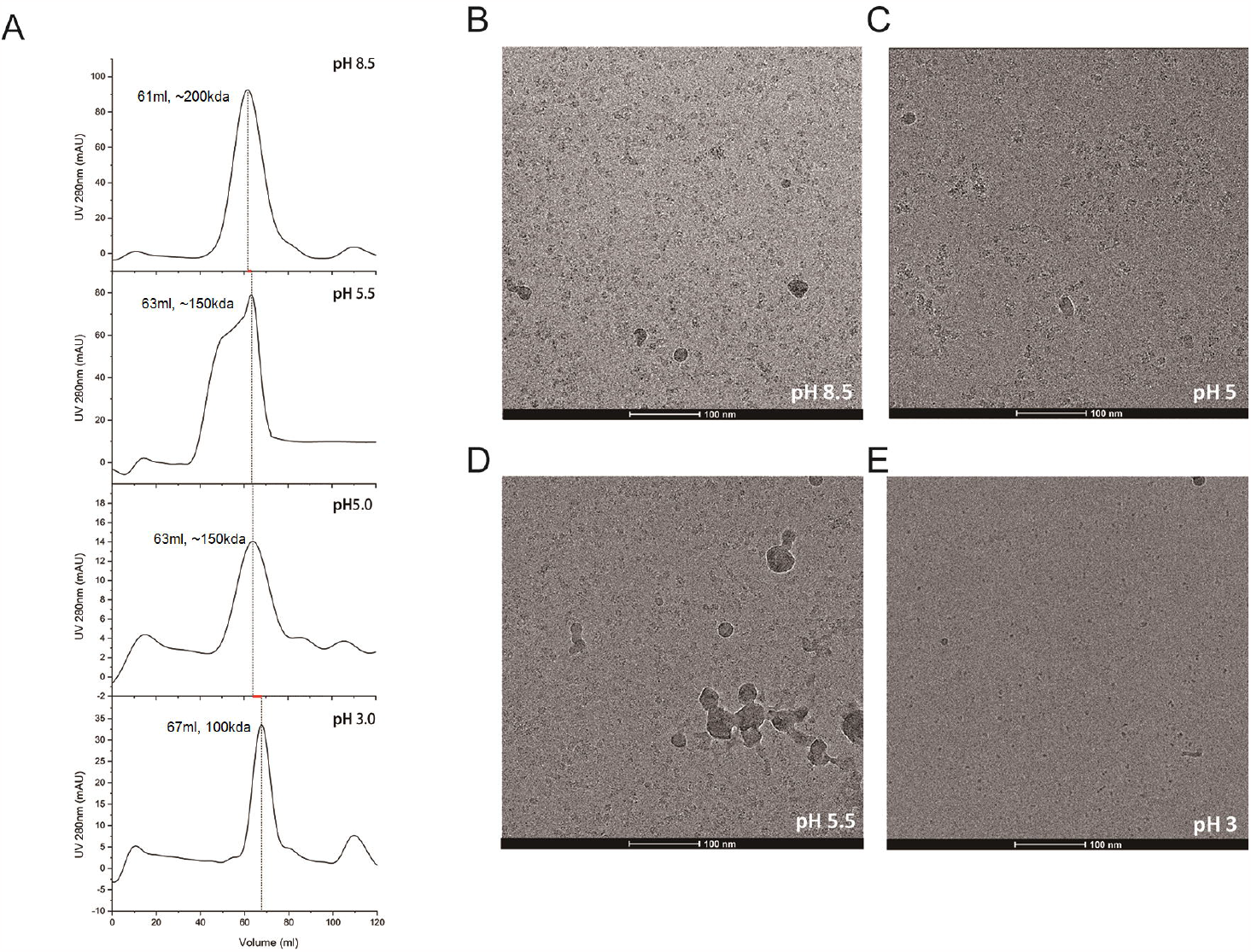
p72 trimer at low pH may undergoes depolymerization. (A) SEC peak position of P72 at different pH. Peak tip positions corresponding to p72 target peaks are marked with dotted lines, and the position differences between dotted lines are represented by red lines. (B-E) CryoEM images of P72 at different pH: pH8.5 (B), pH5.5 (C), pH5 (D), pH3 (E).

The samples at each peak tip were taken to prepare frozen samples, and then checked by cryo-electron microscopy (cryo-EM) at low voltage (Figure 3B). In the cryo-EM photograph of untreated p72 samples, evenly distributed particles with diameters between 7-10nm can be seen. It has been reported that the particle size of p72 trimer should be 8nm in both width and length (16). However, similar particles could not be seen in the cryo-sample of p72 treated under pH 3, only small particles with high contrast could be seen, and the diameter is determined as 3-6nm.

### Cross-linking result

P72 trimer undergoes depolymerization after exposure to acidic environment. Based on this analogy, we examined whether the polymerization state of acid-treated p72 differed from that of untreated sample. To this end, equal amounts of untreated p72 and acid-treated p72 were cross-linked and analyzed by western blot. The result revealed the presence of trimeric p72 in all samples, but the band corresponding to p72 monomer was only shown in acid-treated samples.

Subsequently, the band intensity ratio of p72 monomer to trimer was calculated, which was used to correct for possible loading differences between samples. The result indicated that the amount of p72 trimer in acid treated samples appeared to be reduced along with the reduction of pH (Figure S4).

### Three-dimensional Reconstruction of p72 at pH 5.5

We performed single particle 3D reconstruction of the p72 sample being treated with pH 5.5 acidic environment (Figure 8). A total of 629 .MRC format data files were collected with the pixel size of 1.17Å. Data CTF parameters are determined using CTF Estimation in CryoSparc. A total of 158,530 particles were selected and subjected to 2D classification. After selected 15 2D classes with better quality, ab-initio under C1 symmetry was performed for 3D reconstruction. No specific initial model was provided during the reconstruction process, and the reconstruction result was divided into three classes. Among them, the result of the second class is highly consistent with the p72 trimer. After a round of 3D refinement applying with C3 symmetry, a map of p72 trimer at 4.8Å in resolution was acquired (Figure 8 and 9A). The volume of the trimeric p72 particle was 119.6×10^3^Å^3^ (map contour level=0.3).

As for the third class, another round of 3D reconstruction was carried out. Five reconstruction results were finally obtained, and showed dimer-like particle morphology. The classes with better quality, class 2 and 3 in the 2^nd^ round 3D reconstruction, were optimized and the final resolutions were determined as 14.85Å and 15.91Å (Figure 8). Besides, the particle volume of the p72 treated at pH 5.5 was determined as 85.12×10^3^Å^3^ and 76.53×10^3^Å^3^, respectively (map contour level=0.3) (Figure 8). That is, compared with the p72 trimer, the acid-treated p72 decreased 28.8%-36% in particle size. Combing the result of SEC and single particle 3D reconstruction, we claimed ∼30% reduction in particle volume when p72 trimer was exposed to a low pH buffer (pH 5.5) that mimics the endocytic environment. Correspondingly, two p72 monomers with their F1-G1 loop truncated could be fitted into the electron density (14.85Å in resolution) with CC value ≥0.87. A putative model of the dimeric p72 was proposed based on the superposition result (Figure 4B).

**Figure 4.**
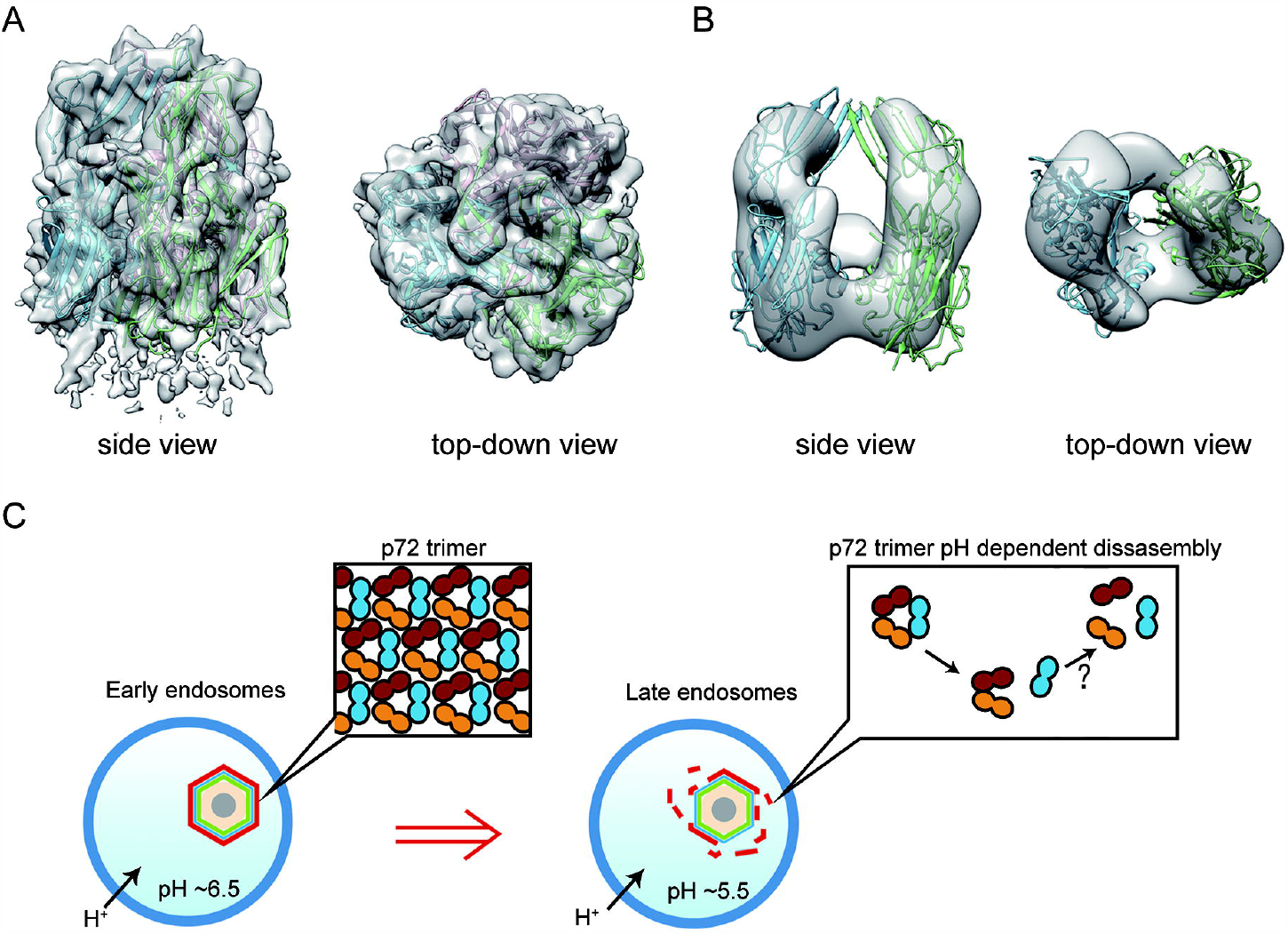
p72 trimer pH-dependent depolymerization may responsible for the rupture of ASFV capsid. (A) Fitting results of the p72 trimer structure (PDB ID: 6KU9) with the electron density at a resolution of 4.8 Å, with a CC value of 0.86. The three monomers are shown in blue, green and pink, respectively. (B) The fitting results of the p72 monomer structure (PDB ID: 6KU9) and the electron density with a resolution of 14.85 Å, the CC value of monomer 1 (blue) and monomer 2 (green) is 0.88 and 0.87, respectively. (C) The putative rupture model of ASFV capsid in the endosomal pathway.

## Discussion

Since the in vitro low pH treatment (pH 5) is sufficient for inducing ASFV disruption, the disassembly does not require other biomolecules provided by the host (7). The disassembly process is likely to depend on at least one pH sensing protein. In this study, we identified the major capsid protein p72 as the pH sensor. It is found that the p72 has the highest histidine content compared with the other structural proteins located on the capsid. Histidine is a pH-sensitive amino acid. Low pH could induce reversible protonation in the imidazole ring of histidine, leading to changes in interactions including hydrogen bond, salt bridge, π stacking and so on, and finally cause conformational changes in protein (13-15). Up to date, the key role of histidine in pH-dependent conformational change has been demonstrated in major histocompatibility complex (MHC) II (17), Hyperpolarization-activated cyclic nucleotide-gated (HCN) (18), Presenilin (19) and influenza virus M2 proteins (20).

Corresponding with the finding that at least 21 histidines are located on the intra-trimer interface, our experiments verified that the p72 trimer undergoes depolymerization under low pH. The results of SEC analysis show that the particle size of recombinant p72 decreases along with lowering pH. Meanwhile, the results of single particle cryo-EM 3D reconstruction showed that only 50% of the particles still maintained the trimeric conformation, and 25% of the particles were found to turn to dimer. In addition, although no structural data were obtained for p72 samples treated with pH 3, the results of size-exclusion chromatography and cryo-electron microscopy strongly support the complete depolymerization of p72 trimer under this condition. This result implies a drastic disassembly process when ASFV infect host by ingestion pathway, as the pH in stomach could be lower than pH 3.

Figure 4C summarizes our proposed an ASFV uncoating model. In the endocytic pathway, due to the influence of low pH and the existence of histidine on the intra-trimer contact surface, the p72 monomers repel each other, a single monomer was dissociated, and lead to the cracking of capsid. The resulting dimer is highly unstable and may be prone to further depolymerization.

It however remains unclear which part of p72 plays the role as a molecular switch in the depolymerization. The dimeric p72 may be highly flexible, and result in the poor resolution. Despite more than 20,000 particles were applied for 3D reconstruction, the final resolution of dimer is still greater than 15Å, and no further structural information can be obtained. According to the histidine distribution analysis, we suggested the interlocked F1-G1 loop in p72 trimer as a candidate. In structural studies on other DJR-containing viruses including PRD1 (21), Faustovirus (22), Adenovirus (23) and Mavirus(24), the interlocked N-terminal base domain is considered relevant for the stability of trimeric MCP. Among them, similar trimer depolymerization has previously been reported in Mavirus virophage (24). Molecular dynamics study on Mavirus suggested that the low pH induced N-terminal conformational changes is responsible for the trimer dissociation (24). However, the ordered N-terminal base domain of p72 trimer can only be observed in ASFV capsid in situ, but is unable to resolve in the recombinant p72 trimer (5, 25, 26). In this study, the ordered N-terminal base domain density of P72 trimer was also not observed. This phenomenon suggests that ordered N-terminal base may not be a necessary element for maintaining the conformation of ASFV p72 trimer.

Besides, the histidine content of ASFV p72 is notably exceeding its homologues from NCLDVs and other dsDNA viruses. The histidine content of ASFV p72 is at least 1.5 folds of that in other NCLDVs MCP. Considering the ASFV is the only enveloped icosahedral NCLDVs that infected mammals, the environmental pressure from the host may result in the capsid disassembly mechanism of ASFV being different from the other NCLDVs.

In summary, this study verified the ASFV major capsid protein p72 as pH sensitive protein, and provided a new insight into the mechanism of capsid disassembly. However, further investigation is still needed to illustrate whether the same trimer depolymerization happens in the acidic full particle.

## Materials and Methods

### Construction and identification of p72 expression vector

To obtain correctly folded P72 trimer, the expression strain was constructed by referring to the method of co-expressing P72 and B602L reported by Qi Liu et al. (26). P72 gene, B602L gene, yeast GAL1 promoter and ADH1 terminator were constructed into the plasmid to form a gene expression box. The gene expression box with homologous recombination arm was amplified by PCR to prepare the repair template. Using CRISPR-Cas9 technology, the gRNA that recognized GGATTTAGGAATCCATAAAA was co-expressed, and the gene expression box was inserted into the Ty2 retrotransposon of multiple copies of saccharomyces cerevisiae by homologous recombination to achieve multiple copy gene expression. The repair template and pCas-ty2 plasmid were co-transformed into yeast cells by electroporation. Then the positive colonies were analyzed by SDS-PAGE.

### Expression, Purification and Identification of Recombinant P72 Protein

Monoclonal saccharomyces cerevisiae colonies were selected and inoculated in 50mL liquid YPD medium and shaken overnight at 30□. The overnight culture products were transferred to large bottles of liquid YPD medium for 10-fold expansion culture, and continued to be cultured at 30□ until OD600 reached 3.0. The culture products were collected and centrifuged at 6000rpm for 10min at 4□ to collect the cell precipitates. After the precipitation was suspended with 50mL washing buffer, yeast cells were disrupted by a high-pressure homogenizer at 4□ and 1800bar pressure. The products were centrifuged at 17000rpm at 4□ for 60min, and the supernatant was collected. The target protein was purified by strep-Tactin XT gravity-flow column.

### Acid Treatment and SEC Analysis of Recombinant P72 Protein

The co-expression of p72-B602L without acid treatment and after acid treatment was analyzed by size exclusion chromatography (SEC). The concentrated purified p72 sample was slowly added to a low pH buffer and incubated at 37□. The homogeneity and aggregation state of untreated and acid-treated p72 were verified by SEC by column chromatography. Care should be taken to avoid inhaling air bubbles during the process and it should be carried out in a subzero low temperature environment. Finally, the collected samples were analyzed by SDS-PAGE.

### Western blot analysis of cross-linked p72 samples

Concentrate the purified p72 trimer to 1mg/ml, slowly add the p72 protein into the low pH buffer (100mM citrate buffer of pH 3, pH5 and pH5.5) according to the ratio of p72:buffer=1:1, blow and mix well. The p72-buffer mixture was acidified at room temperature (25 □) for 30min. After the acidification reaction, adjust the pH of p72-buffer mixture to 6-7 with 0.2M Na_2_HPO_4_.

Based on the above p72 acidified samples, take 19ul of the above p72-buffer samples and mix them with 1ul of 1% glutaraldehyde, incubated at 4 □ for 20min, then place them at room temperature (25 □) for 10min to complete the protein crosslink.

For protein analysis, samples were diluted with 5 × Loading sample buffer, heated at 95°C for 10 min, and loaded on 6% SDS-PAGE gels. Gels were run at 110 V for 80 min in l × Tris-Glycine running buffer and stained with Coomassie Blue Stain solution. Following SDS-PAGE, proteins were electrically transferred onto 0.2 pm nitrocellulose membrane (Pall), 100mA, 140min. The membranes were blocked in 2% skim milk in PBS at 4°C overnight. Primary antibody at a 1:5,000 dilution of anti-strep antibody in 5% skim milk in PBS, was added and incubated for 4 h at room temperature. Membranes were washed with TBST (3 × for 10 min each) and added with secondary antibody 1:5,000 dilution of goat anti-mouse IgG-HRP in 5% skim milk (PBS) for incubation at room temperature for 1.5 h. Membranes were then washed again with TBST (3 × for 10 min each) and developed using TMB (3,3,5,5’-Tetramethylbenzidine, Sigma Aldrich) and hydrogen peroxide for imagining.

### Preparation of Frozen Samples of P72 Protein

Sample was added to a copper net covered with carbon film and quickly immersed in liquid ethane cooled by liquid nitrogen to rapidly form glassy ice. Vitrobot Marker IV of Tsinghua University cryo-electron microscopy platform was then used to prepare samples. After sample preparation, transfer it to a vacuum cup containing liquid nitrogen, and make sample records. Finally, 200keV Arctica transmission electron microscope (Falcon II camera) of Tsinghua university cryo-electron microscope platform was used to examine the frozen samples. According to the observed ice thickness, protein particle contrast and protein concentration, the frozen samples were optimized by adjusting blot parameters and protein concentration.

### Collection and Processing of Cryo-electron Microscopy Data

Data collection of frozen samples in this study was completed on the cryo-EM platform of Tsinghua University. Untreated samples were collected using a Titan3 electron microscope equipped with a Gatan K2 Summit direct electronic counting camera. FEI Talos Arctica was used to collect samples treated with acid. Samples were loaded by Autoloader. AutoEMation2 software is used for automatic data collection. According to the data collection process, each photo location was given a movie stack containing 32 frames, and each set was stored as a. MRCS file. The dosef_logviewer software was used to perform a MotionCorrection on each photo in the collection, resulting in a combined multi-frame photo (*SumCor. mrc), according to which the data collection process can be adjusted at any time.Finally, CryoSparc and RELION software were used for routine processing procedures and data analysis.

