## Supplementary for "African Swine Fever Virus major capsid p72 Trimers function as a pH sensor during uncoating process of virus endocytosis"

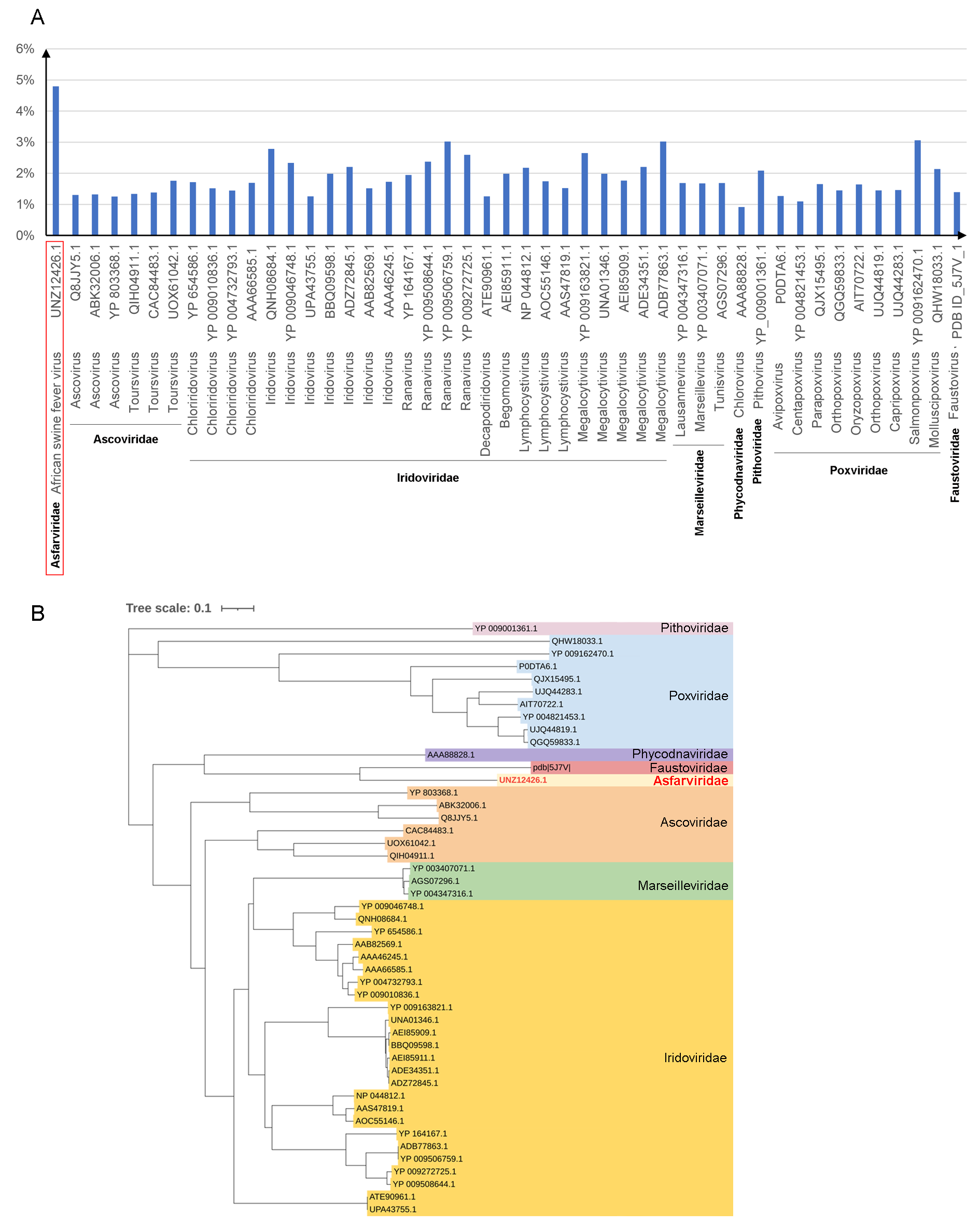

**Figure S1.** Phylogenetic analysis of the NCLDVs MCP

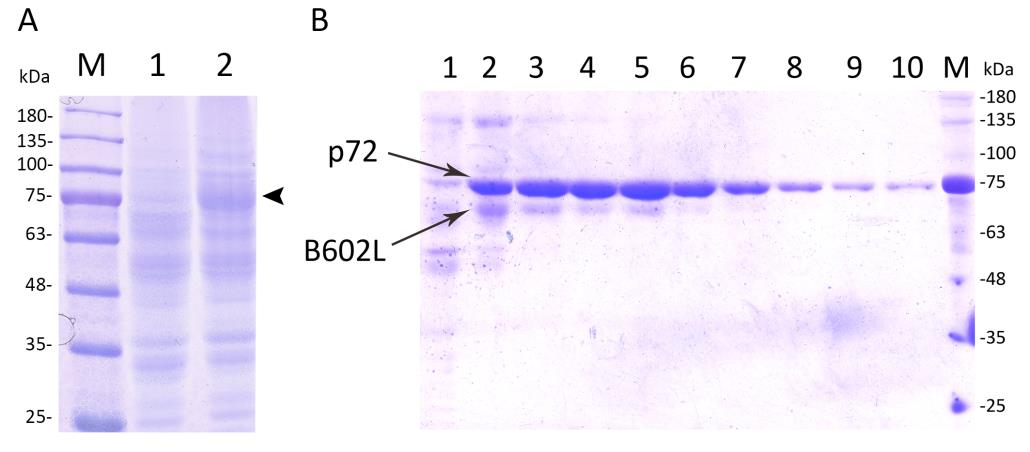

**Figure S2.** SDS-page analysis of the purified product was performed. (A): M: protein marker, 1: control, 2: p72-B602L expressing strain, the location corresponding to the target protein is marked by black arrow. (B): 1-10: the bands corresponding to p72 protein and B602L protein from the 1st to 10th fraction are marked by arrows, M: protein marker.

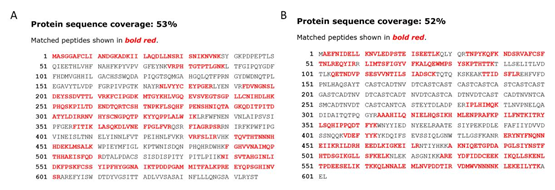

**Figure S3.** MS analysis of protein coverage results. (A): P72 protein. (B): B602L protein.

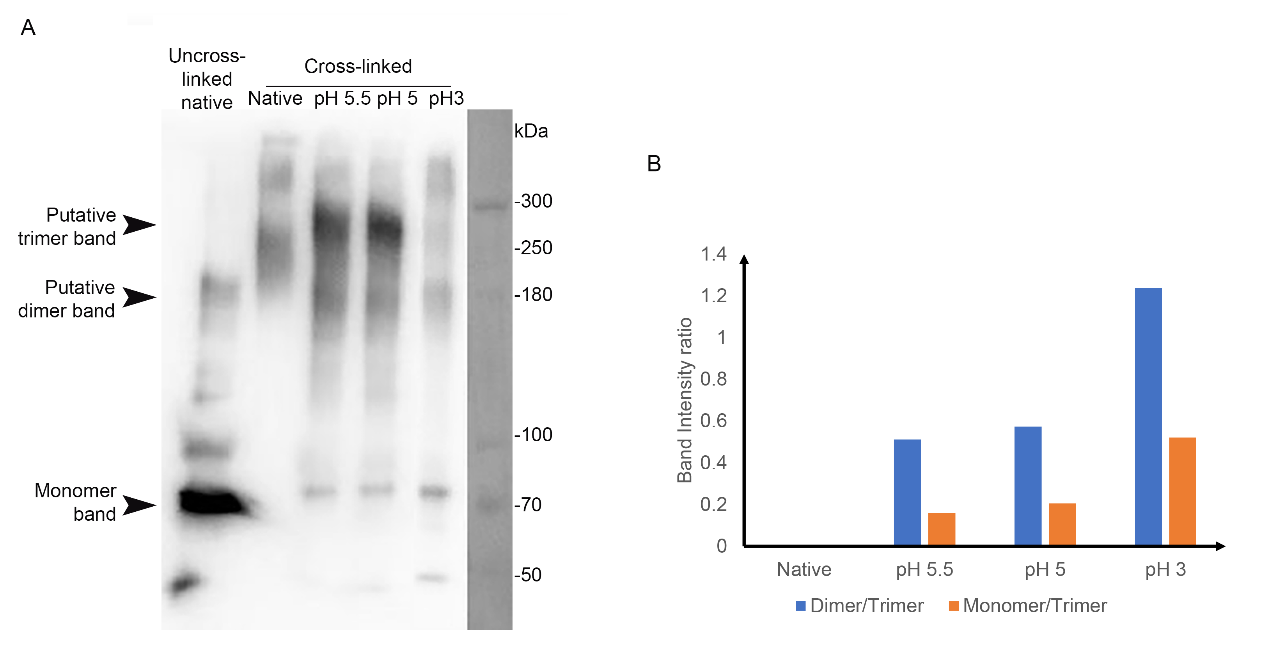

**Figure S4.** (A) Western blot analysis of cross-linked p72 samples. (B) Bands representing trimeric p72 and the monomer in panel A were quantified by GelAnalyzer. By computing the band intensity ratio, differences in sample loading were taken into account.

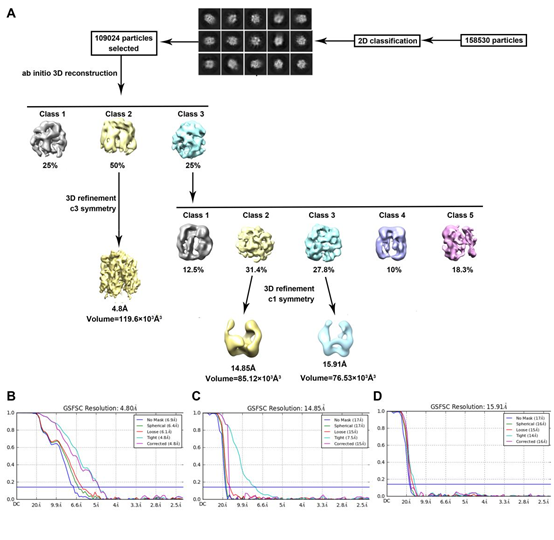

**Figure S5.** S**i**ngle-particle 3D reconstruction of acid-treated p72 samples. (A). In 3d reconstruction, the particle proportion under each classification is shown as a percentage. The resolution of the final optimized reconstruction result was determined by FSC curve (B), (C) and (D), and marked below the final reconstruction result. The particle volume was measured by Chimera at 0.3 contour level and marked below the final reconstruction result.

**Table S1.** The host range of NCLDVs

| **Type of virus** | **Host** | | |
| --- | --- | --- | --- |
|  | **Kingdom** | **Phylum** | **Order/Family** |
| Ascoviridae | Animal | Arthropoda | Lepidoptera（Noctuidae, etc.） |
| Asfarviridae | Animal | Vertebrata Chelicerata | Suidae  Ixodida |
| Iridoviridae | Animal | Vertebrata  Arthropoda | Cypriniformes; Peracarida; Salmoniformes; Cryptobranchidae; Ranidae; Salamandroidea; Siluroidea; Gekkoninae; Serranidae; Sinipercidae  Anophelinae; Oniscidea; Astacidae; Lepidoptera（Noctuoidea, etc.）; Gryllidae |
| Marseilleviridae | Protozoa | Amoebozoa | Amoeba |
| Mimiviridae | Protozoa | Amoebozoa | Amoeba |
| Phycodnaviridae | eumycetes  mycota  Chromalveolata | Chlorophyta  Phaeophyceae  Haptophyta | Chlorellales  Rhodomonas、Ectocarpales、Synedra  Prymnesiales、Isochrysis |
| Pithoviridae | Protozoa | Amoebozoa | Amoeba |
| Poxviridae | Animal | Vertebrata  Arthropoda | Primates、Rodentia、Artiodactyla（Bovinae、Camelidae）、Passeriformes、Carnivora（Canidae、Felinae）、Elephantidae, etc.  Lepidoptera（Noctuoidea, etc.）、Coleoptera |
| Faustoviridae | Protozoa | Amoebozoa | Amoeba |
| Pandoraviridae | Protozoa | Amoebozoa | Amoeba |

^a^ https://www.genome.jp/virushostdb
